# Sex-specific multi-level 3D genome dynamics in the mouse brain

**DOI:** 10.1101/2021.05.03.442383

**Authors:** Devin Rocks, Mamta Shukla, Silvia C. Finnemann, Achyuth Kalluchi, M. Jordan Rowley, Marija Kundakovic

**Affiliations:** Department of Biological Sciences, Fordham University, Bronx, NY, USA; Department of Genetics, Cell Biology and Anatomy, Nebraska Medical Center, Omaha, NE, USA

**Author notes:** Corresponding authors. (M.K.); (M.J.R.). These authors contributed equally to this work (Devin Rocks, Mamta Shukla).

## Abstract

The female mammalian brain exhibits sex-hormone-driven plasticity during the reproductive period. Evidence implicates chromatin dynamics in gene regulation underlying this plasticity. However, whether ovarian hormones impact higher-order chromatin organization in post-mitotic neurons *in vivo* is unknown. Here, we mapped 3D genome of ventral hippocampal neurons across the estrous cycle and by sex in mice. In females, we found cycle-driven dynamism in 3D chromatin organization, including in estrogen-response-elements-enriched X-chromosome compartments, autosomal CTCF loops, and enhancer-promoter interactions. With rising estrogen levels, the female 3D genome becomes more similar to the male genome. Cyclical enhancer-promoter interactions are partially associated with gene expression and enriched for brain disorder-relevant genes. Our study reveals unique 3D genome dynamics in the female brain relevant to female-specific gene regulation, neuroplasticity, and disease risk.

## Introduction

Sex differences in brain physiology and disease result from the interplay of sex hormones and sex chromosome-linked genes (*1*). Brain sexual differentiation is established during perinatal development and the process continues across peripubertal and adult life. From puberty to menopause, cyclical fluctuations in ovarian hormones represent a female-unique experience, and while necessary for reproductive function, they are associated with substantial behavioral and brain plasticity (*2*), and with increased female risk for certain brain disorders such as anxiety and depression (*3*).

Female sex hormones, estrogen and progesterone, have receptors widely distributed in the brain and exert potent neuromodulatory effects (*4, 5*). In rodents, the estrous cycle is characterized by variation in dendritic spine density in the hippocampus, largely driven by fluctuating estrogen levels (*6, 7*). Other aspects of hippocampal physiology, such as long-term potentiation (*8*) and dentate gyrus neurogenesis (*9*), also show cyclic patterns and are enhanced during the high-estrogenic phase of the cycle. Although studies in humans are limited, recent findings indicate that the structure and functional connectivity in the human female brain are comparably dynamic and vary with the menstrual cycle (*10–13*). However, little is still known about molecular mechanisms underlying the sex hormone-induced, dynamic nature of the female brain.

We recently showed that chromatin accessibility, a major mechanism controlling gene expression, varies with the estrous cycle and sex in the ventral hippocampus, a brain region essential for emotion regulation in mice (*14*). We linked these sex-specific chromatin dynamics to changes in neuronal gene expression, neural plasticity, and anxiety-related behavior (*14*). However, whether sex hormones are able to dynamically change the higher-order chromatin organization in post-mitotic neurons of the brain remains unknown. Three-dimensional (3D) genome organization allows interactions of genes with their distant cis-regulatory elements, through chromatin looping and compartmentalization, and is thought to play a major role in transcriptional regulation (*15–17*). Within the brain, 3D genome remodeling has only recently been implicated in neuronal differentiation (*18*) and function (*19, 20*), neuronal activity-dependent gene regulation (*21, 22*), and memory formation (*23–26*), but whether there are sex differences and sex hormone-mediated influences on this regulation remains unknown.

To address this question, we profiled 3D genome organization in adult ventral hippocampal (vHIP) neurons across the estrous cycle and by sex using an unbiased chromatin conformation capture (Hi-C) method combined with DNA fluorescence *in situ* hybridization (FISH) for candidate loci. We integrated 3D genome data with chromatin accessibility (ATAC-seq) and gene expression (RNA-seq) data on the same biological samples. In addition to sex differences, our study shows significant multi-level changes in 3D genome organization across the estrous cycle, which are partially associated with chromatin accessibility and gene expression changes and enriched for brain disorder-relevant genes and pathways. Our study reveals unique 3D genome dynamics in the female brain that has the potential to contribute to both brain and behavioral plasticity and female-specific risks for brain disorders.

## Results

### Study design

For this study, we performed the Hi-C method on vHIP neurons isolated from 11 week-old male and female mice (**Fig. 1A**). To explore the effect of the estrous cycle on 3D genome dynamics in the female brain, we tracked the cycle comprehensively over three consecutive cycles (*14*) and included two female groups in the following estrous cycle stages: proestrus (high estrogen-low progesterone) and early diestrus (low estrogen-high progesterone) (*14*), reminiscent of the human follicular and luteal phase, respectively (**Fig. 1A**). From all three groups, vHIP tissue was first cross-linked to “fix” protein to DNA and preserve 3D interactions, nuclei were isolated, and neuronal (NeuN+) nuclei were purified using fluorescence-activated nuclei sorting (FANS) (*27*). The Hi-C assay proceeded with DNA digestion by restriction enzymes, filling the digested ends and labelling them with biotin, and ligation (**Fig. 1A**). Finally, the proximally-ligated DNA was fragmented, and the biotinylated fragments were enriched and used for Hi-C library preparation (**Fig. 1A**). We performed bioinformatics analysis on Hi-C libraries to explore different levels of 3D chromatin organization including chromosome territories, compartments, CTCF loops, and enhancer-promoter (E-P) interactions (**Fig. 1B**). Hi-C experiments were performed in triplicates which clearly clustered by group (**Fig. 1C**). Considering the high correlation between the replicates in each group, we pooled the data by group for all subsequent analyses resulting in 0.53-0.70 billion sequenced reads per group with 0.36-0.48 billion useable reads after alignment and quality filtering (**Suppl. Table 1**). To explore the role of 3D genome organization in the regulation of gene expression in vHIP neurons by sex and estrous cycle, the Hi-C data were integrated with our chromatin accessibility (ATAC-seq) and gene expression (RNA-seq) data, which were also generated in triplicates (*14*).

**Figure 1.**
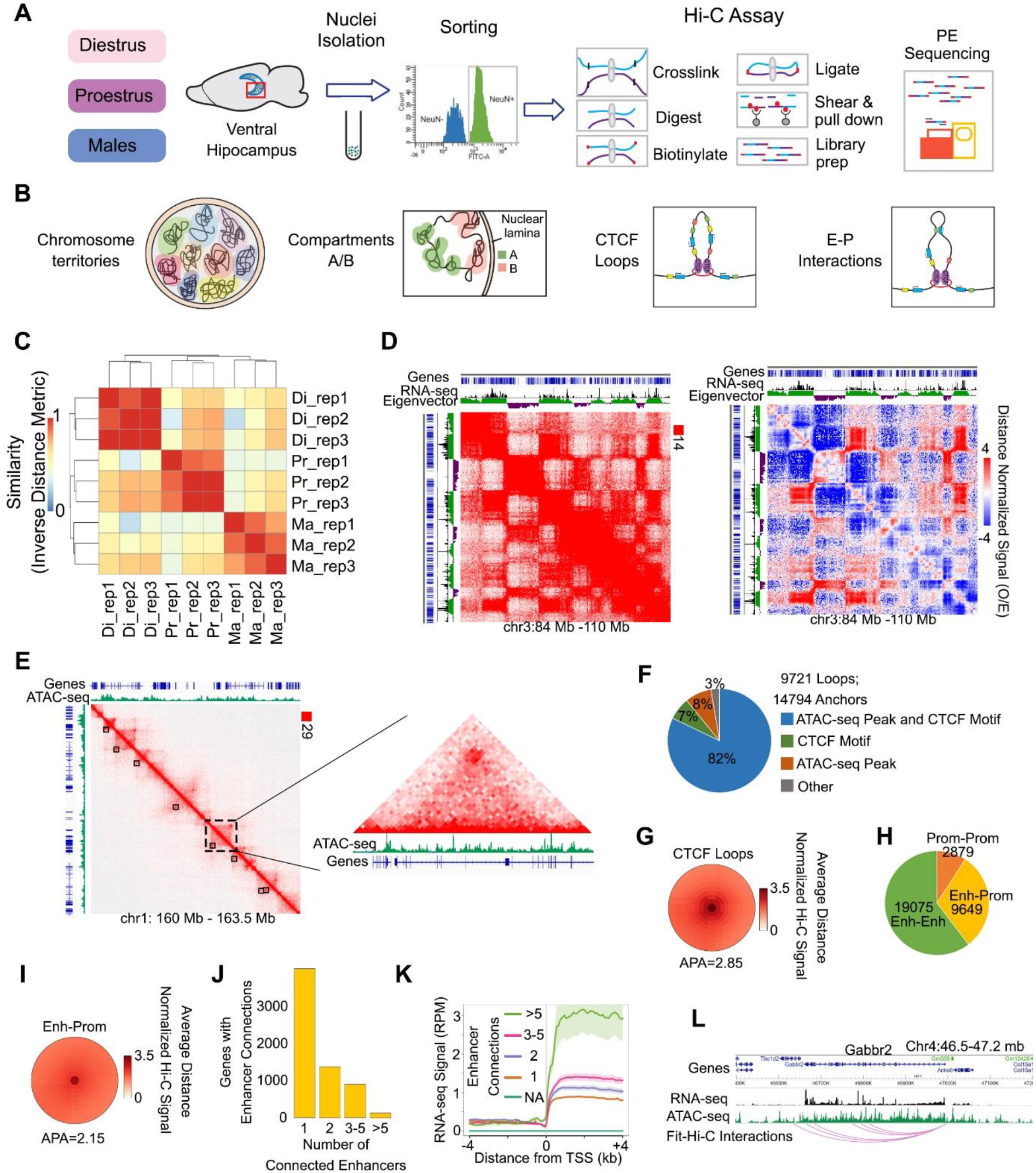
3D genome organization of vHIP neurons across the estrous cycle and sex. The Hi-C assay was performed on sorted neuronal (NeuN+) nuclei isolated from the ventral hippocampus of diestrus, proestrus, and male mice (**A**) to study different levels of 3D chromatin organization (**B**). Biological replicates clustered per group (**C**) and the pooled diestrus group was selected as a representative to explore 3D genome organization. The Hi-C contact matrix (left) and distance normalized matrix (right) are presented with the RNA-seq data overlaid onto the eigenvector signal; the plaid pattern and corresponding gene expression profile indicate the presence of two compartments, active and inactive, called at 25 kb resolution (**D**). The high-intensity punctate signals represent CTCF loops, identified at 10 kb resolution (black squares) and shown with overlapping ATAC-seq signal (**E**), with the majority of loop anchors containing both an ATAC-seq peak and CTCF motif (**F**); the average distance normalized loop signal is shown (**G**). As a third component of chromatin organization, enhancer-promoter (E-P) interactions (**H**) exhibit a weaker distance normalized Hi-C signal (**I**) and the number of these interactions varies significantly across genes (**J**) and correlate with gene expression (**K**). Gabbr2 is represented as an example of a gene with multiple E-P connections and high expression levels (**L**). IGV tracks show merged RNA-seq and ATAC-seq data with the corresponding E-P interactions derived from 3 biological replicates of the diestrus group.

### Ventral hippocampal neurons display known features of 3D genome organization

Across all samples and groups, we found that the majority (72-74%) of Hi-C interactions in vHIP neurons are intra-chromosomal, with only a limited degree of interactions found between chromosomes (26-28%, **Suppl. Table S1**), consistent with the existence of distinct chromosome territories in these neurons (**Fig. 1B**). Using the diestrus group as a representative example, we then explored other features of 3D genome organization including chromosomal compartments, CTCF loops, and enhancer-promoter (E-P) interactions (*16*) in our Hi-C data (**Fig. 1C-L, Fig. S1**). Within each chromosome, the Hi-C contact maps showed a recognizable plaid pattern of interactions, indicating chromatin segregation into two major compartments, A (active) and B (inactive) compartments (*28*) (**Fig. 1B, 1D**), called at 25 kb resolution. These compartments are defined by the eigenvector or first component of a principal component analysis, with positive values defining the A compartment and negative values indicating the B compartment (**Fig. 1D**) (*28*). When Hi-C maps were overlapped with our RNA-seq data, as expected, the A compartment correlated with transcriptionally active genes while the B compartment was generally associated with inactive genes (*28*) (**Fig. 1D, Fig. S1A**).

We next examined high-intensity DNA loops commonly referred to as CTCF loops (**Fig. 1B**) which are visually apparent as strong punctate signals in Hi-C maps (*16*) (**Fig. 1E**). Using Significant Interaction Peak caller (SIP) (*29*), we were able to call a total of 9,721 punctate loops in vHIP neurons at 10 kb resolution (**Fig. 1F-G**). Across 14,794 detected loop anchors (**Fig. 1F**), we confirmed that the top enriched motif was the CTCF motif (**Fig. S1B**). Moreover, when overlapped with ATAC-seq data, the vast majority (82%) of the called loops corresponded with both ATAC-seq peaks and the CTCF motif, while few of them corresponded with either CTCF motif only (7%), ATAC-seq peak only (8%), or neither criteria (3%, **Fig. 1F**).

Finally, we used Fit-Hi-C2 (*30*) with ATAC-seq peaks as a proxy for enhancers to call enhancer-promoter (E-P) interactions in vHIP neurons with 10-kb resolution. This analysis resulted in a total of 19,075 enhancer-enhancer interactions, 9,649 enhancer-promoter interactions, and 2,879 promoter-promoter interactions (**Fig. 1H**). On average, the intensity of the E-P interactions (APA score= 2.15; **Fig. 1I**) was weaker than that of the CTCF loops (APA=2.85; **Fig. 1G**), suggesting that E-P interactions are less stable or more dynamic than CTCF loops. We also found that the number of enhancer connections varied significantly among genes (**Fig. 1J**) and was directly correlated with gene expression level in vHIP neurons (**Fig. 1K**). In particular, the highest RNA-seq signal was detected in genes having numerous (>5) enhancer interactions (**Fig. 1K, Suppl. Table 2**), and these multi-enhancer genes showed an enrichment for pathways important for neuronal function and hormone signaling (**Fig. S1C**). In fact, by exploring the data from the mouse neural differentiation study by Bonev et al. (*18*), we found that the expression of our multi-enhancer genes is likely to emerge with neuronal linage specification (**Fig. S1D-F**). These genes were found to be enriched for repressive H3K27me3 histone marks in embryonic stem (ES) cells while this Polycomb-mediated repression appears to be lost in neuronal precursor cells (NPCs) and cortical neurons (*18*) (CNs, **Fig. S1D-E**). We also found that the multi-enhancer interactions that we identified in our vHIP neurons may be emerging during the acquisition of neuronal fate; these interactions are stronger in NPCs than in ES cells and are further strengthened upon neuronal differentiation (**Fig. S1F**). In general, when compared across tissues, our identified multi-enhancer genes show highest expression in the brain including in the cerebral cortex and hippocampal formation (*31*) (**Fig. S1G**). As an example, a highly expressed, brain-specific gene *Gabbr2*, encoding a subunit of the GABA-B receptor, interacts with multiple (6) putative enhancers located in accessible chromatin regions up to 360 kb downstream of the *Gabbr2* promoter (**Fig. 1L**, **Fig. S1H**).

In summary, we show that the Hi-C data from sorted vHIP neurons display known features of 3D chromatin organization, including chromosomal compartments, CTCF loops, and E-P interactions, which are strongly associated with chromatin accessibility states and gene expression in these cells.

### Compartmental organization of the X chromosome is sex- and estrous cycle-dependent

We next compared the compartmental organization of vHIP neurons across the three groups – diestrus, proestrus, and males (**Fig. 2, Fig S2**). We found no significant difference in the compartmental structure of autosomes in either the male to female (Male vs. Die) or within-female (Pro vs. Die) comparisons (**Fig. S2A-C**). However, we found that the compartmental organization of the X chromosome varies with both sex and the estrous cycle (**Fig. 2A-B, Fig. S2D**). The sex difference in the X chromosome was expected considering that the eigenvector signal for females is a mixed signal from the active (Xa) and inactive (Xi) X chromosome, while the same signal in males originates from a single (and active) X chromosome (**Fig. 2A**). Although the difference in X chromosome eigenvector in the male to female (Male vs. Die) comparison was more profound than that in the within female (Pro vs. Die) comparison (**Fig. 2A**), differences were significant in both comparisons in terms of both the overall signal intensity (**Fig. 2B**) and the number of bins being affected (**Fig. S2D**). In the proestrus to diestrus comparison, 1,081 bins or 15.9% (27 Mb) of the X chromosome showed changes in compartmental signal (**Fig. S2D**). These differences in compartmental signal were further associated with differences in chromatin accessibility (**Fig. S2E**).

**Figure 2.**
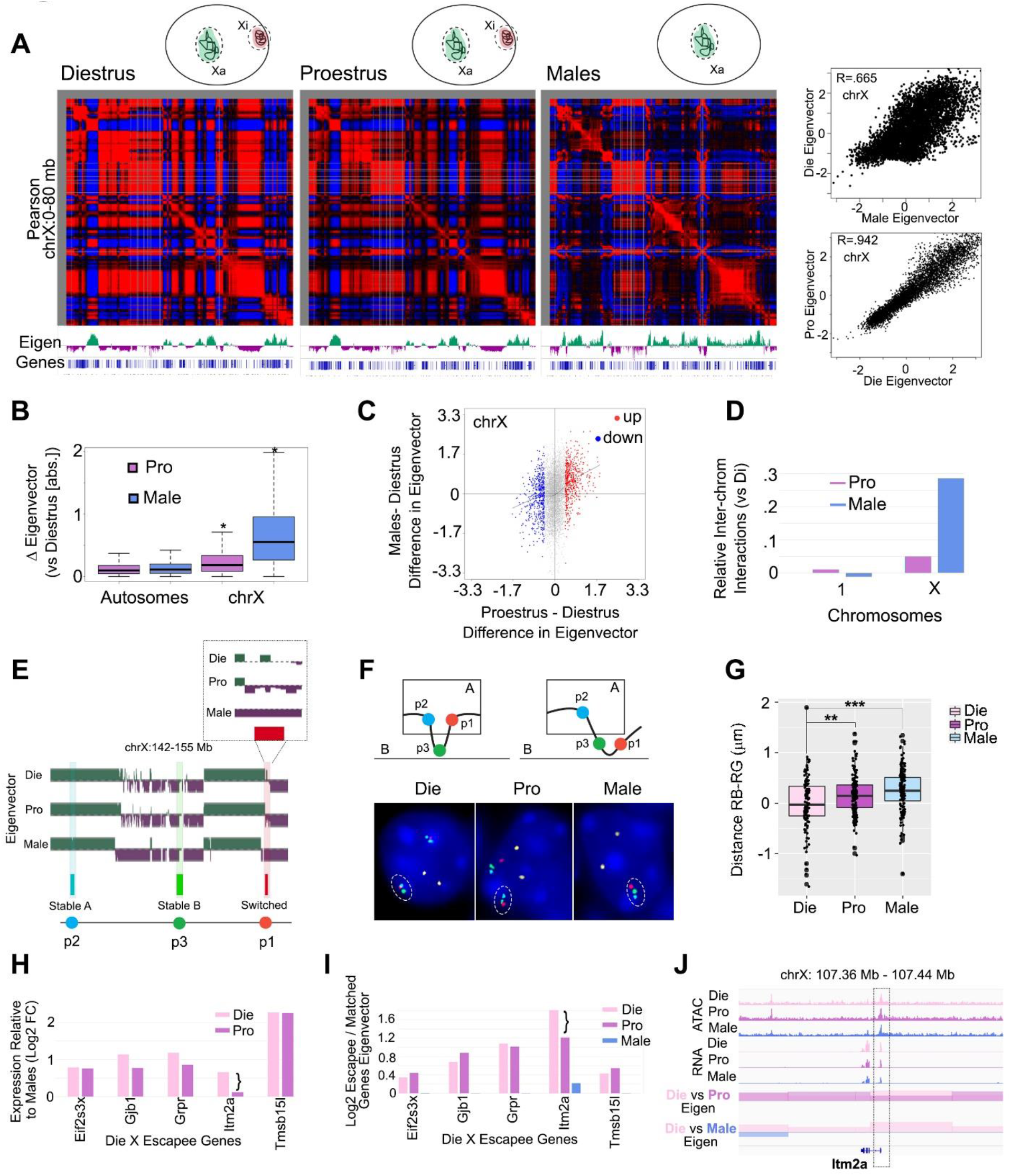
Compartmental organization of the X chromosome in vHIP neurons across the estrous cycle and sex. The correlation matrix and the corresponding eigenvector signal (Eigen) show compartmental profiles along an 80 Mb-segment of the chromosome X in diestrus, proestrus, and male groups (**A**). Females have an active (Xa) and an inactive (Xi) X chromosome while males have only one active (Xa) chromosome (shown in inset), which is consistent with the largely different eigenvector signal in the Die-Male comparison (R=0.665) and more similar profile in the Pro-Die comparison (R=0.942, panels on the right). The eigenvector signal is significantly different in the X chromosome but not in autosomes in both Pro-Die and Male-Die comparisons (**B**). Proestrus and male signals go in the same direction when compared to the diestrus signal (**C**). X chromosome, but not chromosome 1, shows more interactions with other chromosomes in males and proestrus compared to diestrus (**D**). The compartmental profile of a smaller 13-Mb segment of X chromosome is depicted in Die, Pro, and Male, and the location of the designed FISH probes (p1-p3) is shown (**E**) and the expected compartmental change of the region surrounding probe 1, a switch from compartment A to B in Die vs. Pro, was confirmed with FISH (representative images are shown; note: yellow signal is a negative control on chromosome 1) (**F**). FISH data were calculated as the RB-RG distance by subtracting the distance between the center of the red and green signals (RG) from that of the red and blue signals (RB). The analysis was restricted to X chromosomes positive for all three probe signals (n=116-163/group); one-way ANOVA with post-hoc Tukey; **, P <0.01; ***, P <0.001 (**G**). The expression (**H**) and relative eigenvector signal (**I**) of the top five X escapee genes was shown in diestrus and proestrus females compared to males. *Itm2a* gene is shown as an example of X escapee that changes expression and compartmental signal across the estrous cycle (**J**). IGV tracks show merged ATAC-seq and RNA-seq data for diestrus (Die), proestrus (Pro), and males (Male) derived from 3 biological replicates for each group. Die, pink; Pro, purple; Male, blue.

Interestingly, when comparing Pro vs. Die, we also found that the differences in compartmental signal were often going in the same direction as Male vs. Die differences (**Fig. 2C)**, suggesting that the X chromosome compartmental signal in the high-estrogenic proestrus female group is more similar to that of males, when compared to the low-estrogenic diestrus group. Furthermore, we found that the X chromosome in males and proestrus females showed more inter-chromosomal interactions than the X chromosome in the diestrus group (**Fig. 2D**). All described similarities between proestrus and males were specific to the X chromosome and were not found in autosomes (**Fig. 2B, S2F-G**), implying that proestrus may be associated with a higher volume or partial decondensation of the Xi chromosome, making the X chromosome compartmental signal in proestrus more similar to that in males.

To confirm the observed compartmental change in the X chromosome across the estrous cycle, we performed four-color FISH. Based on our Hi-C data, we designed a control DNA probe (p4) located in chromosome 1, and three DNA probes located in the X chromosome: the probes p2 and p3 were in the compartments A and B, respectively, in all three groups; and, the probe p1 was found to be in the compartment B in males while it showed a “switch” between compartments B (Pro) and A (Die) during the estrous cycle in females (**Fig. 2E-F**). We quantified our FISH data by subtracting the physical distance between probes p1 and p3 from the distance between the probes p1 and p2 (**Fig. 2F-G**). We found that p1 and p2 are more physically separated in the proestrus and male samples than in the diestrus samples (**Fig. 2F-G**), confirming the altered compartmental profiles derived from the Hi-C method (**Fig. 2E**).

We further explored whether gene expression changes may be associated with the observed sex- and estrous cycle-dependent compartmental differences. Naturally, we first looked into X chromosome escapee genes which are able to escape the inactivation of the Xi chromosome and are, thus, more highly expressed in females than in males (*32*). Taking the top 5 escapee genes from the Die-Male comparison (**Fig. 2H, Fig. S3A**), we indeed found that these genes are always located in the A compartment in diestrus (**Fig. S3B**) and show a higher eigenvector A compartment signal compared to expression matched non-escapee genes (**Fig. 2I, Fig. S3C-D**). In males, these genes did not show heightened A compartment signal compared to other genes (**Fig. 2I, Fig. S3C-D**). Four of these genes (*Eif2s3x*, *Gjb1*, *Grpr*, and *Tmsb15l*) also showed a higher expression and heightened eigenvector signal in proestrus females compared to males (**Fig. 3H-I**). However, *Itm2a* expression during proestrus was reduced compared to diestrus; this was associated with a decreased eigenvector signal in the Pro-Die comparison making the signal of the proestrus female group, again, more similar to that of males (**Fig. 2H-J**). We also checked the *Htr2c* gene which is located nearby the FISH probes that we designed (**Fig. 2E**) and is another gene that is differentially expressed in the Die-Male but not in the Pro-Male comparison (*14*) (**Fig. S3E**). Importantly, this gene, too, showed a change in the signal between diestrus and proestrus groups, with the proestrus signal being more similar to males, particularly in the area adjacent to the TSS and the end of the gene (**Fig. S3E**).

**Figure 3.**
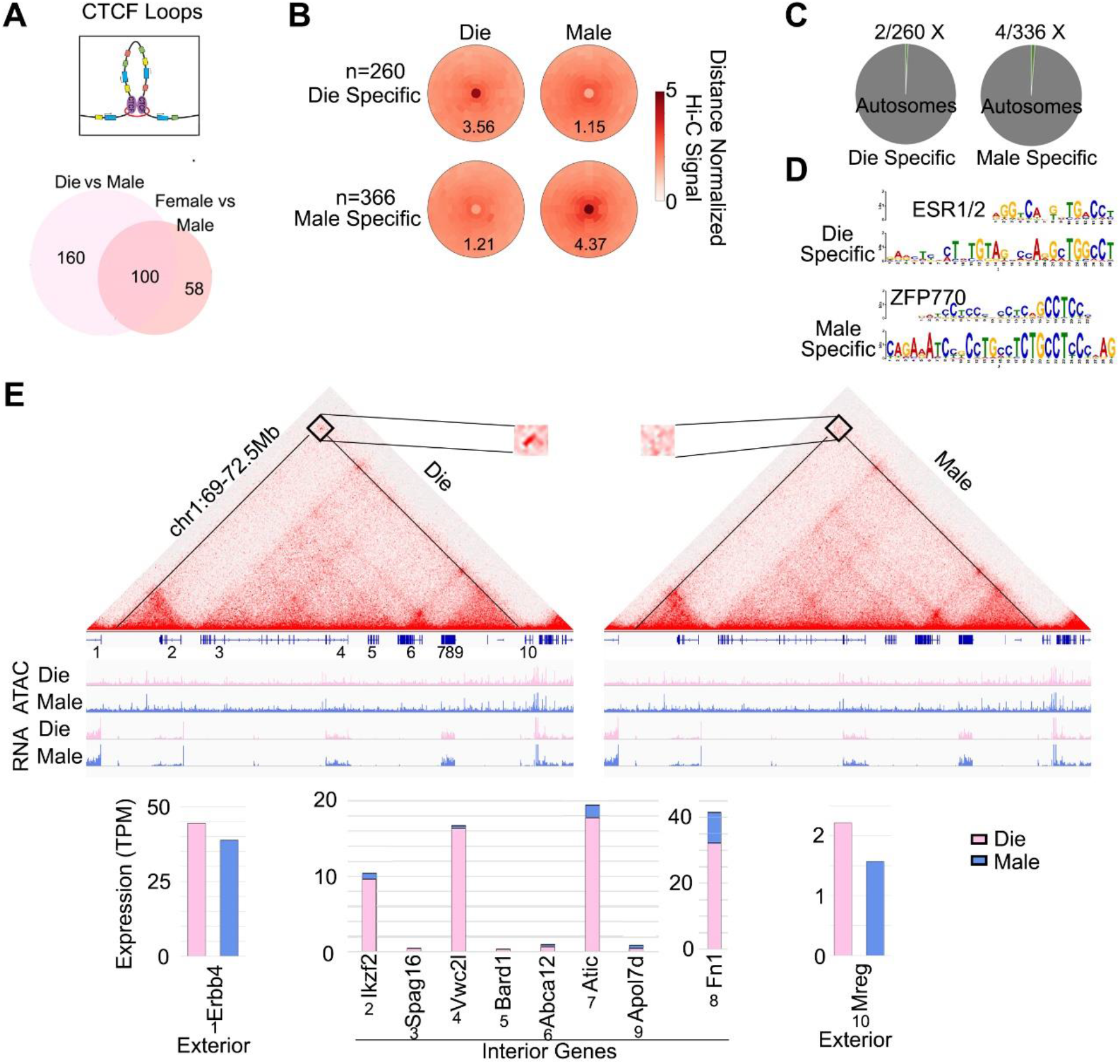
Sex-specific CTCF loops in vHIP neurons. More sex-specific loops can be called in the Die-Male than in the Mixed Female-Male comparison (**A**). Both Die- and Male-specific loops (**B**) are primarily located on autosomes (**C**). Die-specific loops are enriched for the ERα (ESR1) motif while Male-specific loops are enriched for ZFP770 motif (**D**). Presented is a 3.5 MB loop encompassing eight interior genes and flanked by two exterior genes (**E**). IGV tracks show merged ATAC-seq and RNA-seq data for diestrus (Die) and males (Male) derived from 3 biological replicates for each group. Lower panel shows gene expression for each exterior and interior gene. Die, pink; Male, blue.

Finally, to explore which upstream regulators may be driving the observed compartmental differences, we performed the motif analysis of the genomic areas showing differential compartmental signal between the sexes and across the estrous cycle. Interestingly, we found the sex-determining region Y gene (Sry) and Egr2 binding sites to be enriched in the Die-Male comparison (**Fig. S3F**). In the Die-Pro comparison, we found binding sites for the transcription factor Pou3f2 as well as for the response elements for estrogen receptor alpha (ERα, ESR1 motif) and estrogen-related receptor 2 (ERR2, **Fig. S3G**), consistent with hormonal regulation of the higher order chromatin organization in vHIP neurons across the estrous cycle in females.

In summary, we found a significant compartmental change in estrogen response elements-enriched X chromosome genomic areas during the estrous cycle, with proestrus females showing a compartmental profile more similar to males, which was partially associated with gene expression.

### Neuronal CTCF loops vary with sex and the estrous cycle

We next examined the effect of sex and the estrous cycle stage on the CTCF loops in vHIP neurons (**Figs. 3-4**). For the between-sex comparison, we explored comparing either the diestrus group to males or the mixed female group (merged diestrus and proestrus) to males. We found an increased ability (1.65 times) to call sex-specific loops when comparing diestrus to males, as opposed to comparing mixed females to males (**Fig. 3A**), indicating that separating females by the estrous cycle stage helps identify sex differences in chromatin looping. Out of 9721 called loops, we found 260 loops to be stronger in diestrus than in males (Die-specific) and 366 loops to be stronger in males than in diestrus females (Male-specific, **Fig. 3B**). Importantly, unlike compartmental differences, only 6 of these sex-specific loops were on X chromosomes and all others were on autosomes (**Fig. 3C**). Motif analysis found estrogen response elements (the ESR1/2 motif) to be enriched at the diestrus-specific loop anchors while the ZFP770 motif was enriched at the male-specific loop anchors (**Fig. 3D**). Finally, we explored whether gene expression differences were associated with sex-specific loops. We closely examined a diestrus-specific loop encompassing 8 genes and 3.5 Mb at chromosome 1 (**Fig. 3E**). Interestingly, while we found little difference in the expression of the genes interior to the loop, there were more profound differences in the expression of two genes outside of the CTCF loop anchors, including Erbb4 (**Fig. 3E**), the gene encoding the neuregulin receptor. However, while specific loops were associated with transcriptional differences, in general, there was little correlation between differential loops and gene expression (**Fig. S4A**). This finding is consistent with recent evidence suggesting loops are likely only a small component of gene expression control (*33, 34*).

We further explored differential CTCF loops in the within-female comparison and found roughly the same number of differential loops in the Die-Pro comparison (**Fig. 4A**) as we found in the Die-Male comparison (**Fig. 3B**). Located primarily on autosomes (**Fig. 4B**), 370 loops were stronger in diestrus than in proestrus (Die-specific) and 264 loops were stronger in proestrus than in diestrus (Pro-specific, **Fig. 4A**). Interestingly, when compared to diestrus, the loop signal changes in proestrus were largely in the same direction as the loop signal changes in males (**Fig. 4C**). In addition, the motif analysis found estrogen response elements (the ESR1/2 motif) to be enriched in both diestrus-specific and proestrus-specific loops (**Fig. 4D**). Using confocal microscopy, we confirmed that the estrogen receptor alpha (ERα) localizes to the nucleus of vHIP neurons (**Fig. 4E, Fig. S4B**), consistent with its role as a transcription factor and possible regulator of chromatin loops.

**Figure 4.**
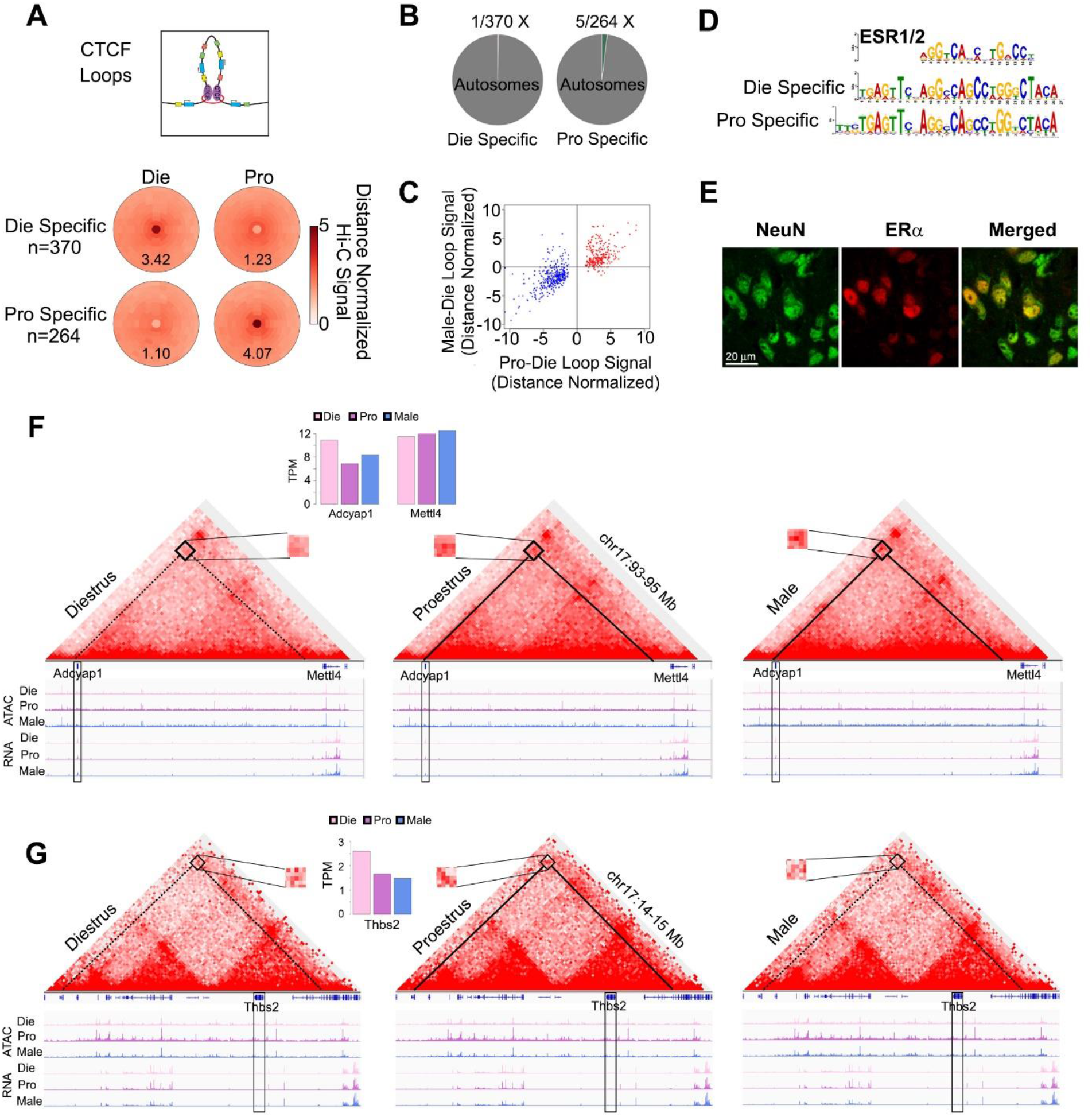
Estrous cycle-dependent CTCF loops in vHIP neurons. Estrous cycle stage-dependent, Die- and Pro-specific loops (**A**) are primarily located on autosomes (**B**); Pro-Die loop signal goes in the same direction as Male-Die signal (**C**). Die- and Pro-specific loops are enriched for the ERα (ESR1) motif (**D**). ERα localizes to NeuN+ nuclei in the ventral hippocampus (**E**, representative proestrus sample is shown). Two example genes *Adcyap1* (**F**) and *Thbs2* (**G**) are presented. Differential loops are depicted with either a solid line (strong loop) or dashed line (weak loop). IGV tracks show merged ATAC-seq and RNA-seq data for diestrus (Die), proestrus (Pro), and males (Male) derived from 3 biological replicates for each group. Insets show gene expression for *Adcyap1, Mettl4* (neighboring gene), and *Thbs2*. Die, pink; Pro, purple; Male, blue.

Similar to what we found for sex-specific loops, there was, in general, little correlation between differential Pro-Die loops and gene expression (**Fig. S4C**). However, specific loops were associated with transcriptional differences. An interesting example is a sex-specific and estrous cycle-dependent loop involving *Adcyap1*, an important stress- and estrogen-sensitive gene implicated in anxiety (*35, 36*) (**Fig. 4F**). This 2Mb loop connects *Adcyap1* and an upstream region of *Mettl4*, is stronger in proestrus and males than in diestrus, and corresponds to differential *Adcyap1* expression among the three groups (**Fig. 4F, Suppl. Table S3**). Notably, another CTCF loop that connects *Adcyap1* and *Mettl4* is present in all groups and may be more important for regulating *Mettl4* expression which shows no expression difference across the groups (**Fig. 4F**). It is worth noting that the observed differential loop also appears in the mixed-female (Die+Pro) to male comparison (**Fig. S4D**), further highlighting that the sex-specific dynamism that we see with proestrus becoming more similar to the male *Adcyap1* gene looping (and the associated gene expression change) is only discoverable if the assay resolution is increased by monitoring the estrous cycle stage (**Fig. 3A**). Another example includes a loop surrounding *Thbs2*, which similar to *Adcyap1*, is also an anxiety-related gene (*37*) that has an estrogen response element in the promoter and can be regulated by ERα (*38*). While *Thbs2* is more highly expressed in diestrus in comparison to both proestrus and males, we found a proestrus-specific loop (**Fig. 4G, Suppl. Table S3**) that is likely to be regulated in a sex-specific way, by varying sex hormone levels in females.

In conclusion, we found sex- and estrous cycle specific CTCF loops in vHIP neurons that are partially associated with gene expression changes and are likely to be regulated by estrogen receptors in females.

### Neuronal enhancer-promoter interactions vary with sex and estrous cycle

Finally, we explored differential E-P interactions and, strikingly, found around 2,000 differential E-P interactions in both Die-Male and Die-Pro comparisons (**Fig. 5A**). Again, when compared to diestrus, the proestrus E-P signal was largely going in the same direction as the male E-P signal (**Fig. 5B**). We found estrogen response elements (the ESR1 motif) to be enriched in diestrus-specific and proestrus-specific EP interactions (**Fig. 5C**), and these elements also appeared as top motifs in female-to-male differential EP interactions (**Fig. S5A**). Although, in general, EP differences did not show significant correlation with gene expression neither in Die-Pro (**Fig. S5B**) nor in Die-Male (**Fig. S5C**) comparisons, around 10% of genes whose expression varies with the estrous cycle (*14*) showed concomitant, cycle-driven differential E-P interactions (**Suppl Table. S3**). For instance, the *Pou3f2* gene, an important psychiatric risk-related gene encoding a brain-specific transcription factor, showed clear estrous cycle-dependent (**Fig. 5D**) and sex-specific (**Fig. S5D**) E-P interaction profiles which were associated with differential gene expression. We further looked into several additional genes with cyclical expression and important neuronal function including *Trhde* (encoding a thyrotropin-releasing hormone degrading enzyme), *Gfra2* (encoding a GDNF family receptor), *Kcnh5* (encoding a potassium voltage-gated channel), *Gsx2* (encoding the GS homeobox 2), and *Zfp580* (encoding the Zing finger protein 580, **Fig. 5E-I**). While some of these genes showed numerous E-P interactions (e.g. *Gfra2*, **Fig. 5F**) and others showed few of them (e.g. *Gsx2*, **Fig. 5H**), the number of which, again, appeared to be associated with the gene expression level, differential interactions were typically associated with the distant genomic regions and correlated with transcriptional differences in an estrous cycle-dependent (**Fig. 5E-I)** and sex-specific manner (**Fig. S6**).

**Figure 5.**
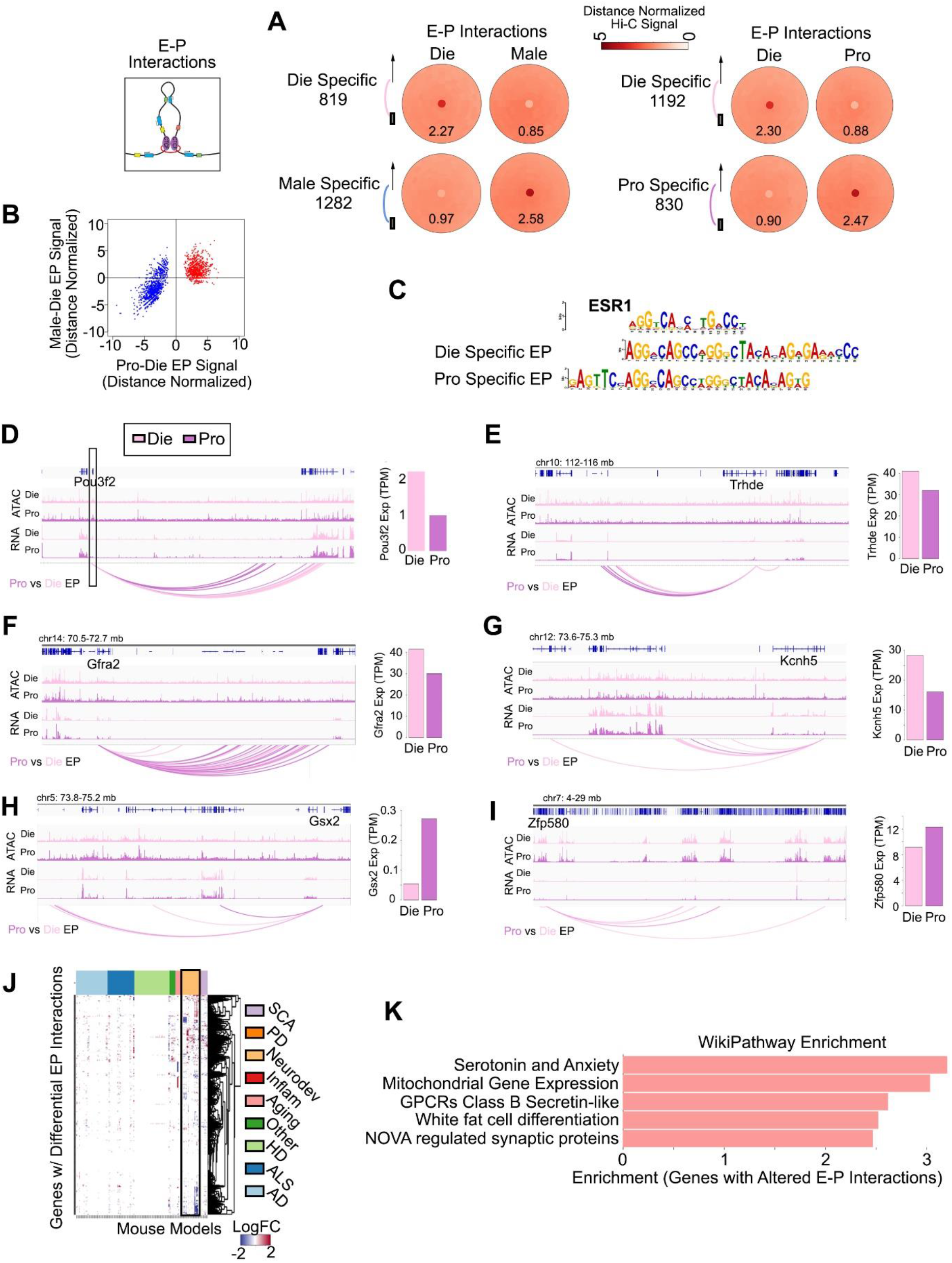
Dynamics of E-P interactions in vHIP neurons across the estrous cycle. The number of differential E-P interactions across sex and the estrous cycle is comparable (**A**) and the E-P signal goes in the same direction in the Pro-Die and Male-Die comparisons (**B**). Across the estrous cycle, in differential Die- and Pro-specific E-P interactions, there is an enrichment of binding sites for ERα (ESR1 motif, **C**). Example genes showing differential E-P interaction profiles and differential gene expression across the estrous cycle include *Pou3f2* (**D**), *Trhde* (**E**), *Gfra2* (**F**), *Kcnh5* (**G**), *Gsx2* (**H**), and *Zfp580* (**I**). IGV tracks show merged ATAC-seq and RNA-seq data for diestrus (Die) and proestrus (Pro) derived from 3 biological replicates for each group. Bar graphs (on the right) show gene expression for each gene. Cyclical E-P interactions are enriched for genes altered in mouse models relevant to neurological disorders (**J**) and several gene pathways including “Serotonin and Anxiety” as the top pathway (**K**).

Finally, we examined which genes and pathways are enriched within estrous cycle- and sex-specific E-P interactions, irrespective of gene expression profiles (**Fig. 5J-K**). We first used a mouse RNA-seq portal that includes mouse transcriptomic datasets related to multiple neurological disorders and aging models (*39*). Interestingly, many of the genes with differential E-P interactions across the estrous cycle, while not showing gene expression changes in our physiological mouse model, exhibit aberrant expression in mouse models of neurological diseases (**Fig. 5J**). In addition, the top pathway shown to be enriched in the diestrus-proestrus comparison was the serotonin and anxiety pathway, further suggesting that the genes relevant to serotonergic function and anxiety-related behavior overwhelmingly show 3D genome organizational changes across the estrous cycle (**Fig. 5K, Fig. S7A**). This pathway was specific to within-female comparison, as we found a more generic G-protein-coupled receptors pathway to be the top enriched pathway in the between-sex (Die-Male) comparison (**Fig. S7B**).

In summary, we found thousands of estrous cycle-dependent and sex-specific E-P interactions that are partially associated with gene expression differences and are enriched for brain disease-relevant genes and pathways.

## Discussion

3D genome organization allows orderly interactions of physically-distant parts of the genome and is thought to play a critical role in gene regulation and cellular function across organs and disease states (*15, 17*). 3D genome remodeling has only recently been implicated in brain development (*18*) and function (*19, 20*), neuronal activity-dependent gene regulation (*21, 22*), and memory formation (*23–26*). However, previous 3D genome studies of the brain were either focused on the male brain or did not explore sex differences. Here we provide the first evidence that 3D chromatin structure in the brain differs between males and females and undergoes dynamic remodeling during the female estrous cycle.

Among different levels of chromatin organization, the compartmental organization of chromosomes is thought to be the most stable one in fully differentiated cells such as post-mitotic neurons (*40*). Here we found no sex- or estrous cycle-driven difference in autosomes, an expected sex difference in the X chromosome, and a surprisingly high degree of X chromosome compartmental dynamics across the estrous cycle. A body of evidence shows that the Xi chromosome in females has a condensed 3D structure and is associated with the nuclear lamina, while the Xa chromosome is more centrally-located in the nucleus, open, and active (*41*). Using Hi-C and FISH as two complimentary approaches to study 3D genome organization (*42*), we show that, with naturally rising estrogen levels in female mice, the X chromosome changes compartmental organization, acquires an interaction profile more similar to males, and is likely to undergo a decondensation event and increase in volume during proestrus. While we observe associated gene expression changes, particularly in X escapee genes, the significance and possible implications of hormonally-driven X chromosome dynamics may go well beyond the regulation of X chromosome-linked genes. For instance, it was proposed that the Xi chromosome may be affecting autosomal gene expression by sequestering heterochromatin factors in the cell (*43*), thus chromatin changes in Xi may affect gene regulation more globally. It is also worth noting that X chromosome decondensation was reported in estrogen-dependent breast cancers (*44*), in which case the observed deleterious X chromosome-reactivation (*45*) could be an extreme case of what estrogen does under physiological conditions. In addition, the variability of X chromosome inactivation was reported to be higher than expected (*46*), including spatially- and temporally-variable escapee genes, which at least in part may be due to the natural variation in hormonal levels driving gene regulation. Together, we anticipate that physiological dynamics in the X chromosome may include global changes in cellular environment affecting neuronal gene regulation and function and this warrants further investigation as it may have important implications for female health and disease.

In general, we observe that estrous cycle- and sex-driven 3D genome changes at all levels of organization are associated with changes in expression of relevant genes. For instance, we see cycle- and sex-dependent compartmental and gene expression changes in the gene encoding the serotonin receptor 2c (*Htr2c*). This receptor is known to play an important role in anxiety-related behavior (*47*), and these data are consistent with our previous findings that diestrus females exhibit higher anxiety indices compared to proestrus females and males (*14*). Furthermore, changes in CTCF looping around genes such as *Adcyap1* and *Thbs2* are associated with sex- and estrous cycle-dependent gene expression differences and further provide a possible link between 3D genome changes and anxiety-related behavioral phenotypes. In particular, *Adcyap1* encodes pituitary adenylate cyclase activating polypeptide (PACAP), a critical sex-specific and estrogen-dependent regulator of the stress response (*48*) which has been associated with sex-specific anxiety-related behavior in rodents (*35*) and female-specific risk for PTSD in humans (*36*). Moreover, we observed differential E-P interactions and changes in the expression of genes such as *Pou3f2* (*49*), *Trhde* (*50*), and *Gfra2* (*51*), which are associated with psychiatric disorders in humans and anxiety and depression-related behaviors in mice. Therefore, considering the importance of 3D genome organization for gene regulation and cellular function, our results provide a new molecular mechanism for sex specific- and sex hormone-driven neuronal gene regulation, neural plasticity, and behavioral phenotypes.

When considering possible mechanisms that drive 3D genome changes in a sex specific- and estrous cycle-dependent manner, it is striking that we found the consistent enrichment of estrogen response elements across all levels of analysis including compartments, CTCF loops, and E-P interactions. Estrogen regulates genes through either classic nuclear estrogen signaling or through membrane-bound estrogen receptors (ERs) (*4*). Our previous findings from the ATAC-seq analysis implicated a membrane-bound ER-mediated mechanism and Egr1 transcription factor in the regulation of chromatin accessibility across the estrous cycle (*14*). However, current Hi-C data strongly imply that 3D genome organization in vHIP neurons in females is driven by nuclear ERα. We know that both estrogen levels and ERα expression vary with the estrous cycle (*14, 52*) and this is likely reflected in the differential, cycle-dependent ERα binding to the genome, thus providing a likely mechanism for changing 3D chromatin organization with varying sex hormone levels. Indeed, studies in breast cancer cell lines showed that the binding of ERs has an instructive role in chromatin looping (*53*) and have proposed a role for steroid receptors as genome organizers at a local and global scale (*54*). Consistent with this, Achinger-Kawecka et al. found that 3D genome remodeling during the development of endocrine resistance in ER+ breast cancer cells is mediated by ERα binding (*55*), further linking ERα with 3D genome organization.

Importantly, we also see a lot of changes in 3D genome interactions that are not associated with gene expression changes in our physiological model. These “non-functional” 3D organizational changes, therefore, may represent an ‘epigenetic priming event’ or could be part of cellular memory associated with cycling events. For instance, in studies of mice with disruptions in the CTCF loop organizer, it was shown that memory-relevant genes do not exhibit hippocampal gene expression changes under basal conditions but are affected in activity-dependent way (*24*). Other studies also showed learning (*56*)- and immunological memory (*57*)-related “epigenetic priming” where epigenomic changes precede changes in gene expression which require another stimulus to be expressed. This molecular priming is consistent with female-specific, reproductive-related physiology where many events across the ovarian cycle including ovarian changes, thickening of the uterus wall, and likely brain structural changes are preparatory rather than functionally relevant events. Strikingly, we also see that estrous cycle-dependent (but not sex-specific) differential E-P interactions are enriched for neurological disorder- and serotonin and anxiety-relevant genes. Both clinical and basic studies show that the increased female risk for depression and anxiety disorders is associated with fluctuating sex hormone levels (*3*). While varying sex hormones by themselves may not be sufficient to trigger the disorder, they may “prime” the female brain for increased vulnerability that can be precipitated by stress and other risk factors. Therefore, the molecular priming effects that we see in the form of 3D chromatin organizational changes across the ovarian cycle may represent the molecular basis for female-specific vulnerability for certain brain disorders such as anxiety and depression.

In summary, our study reveals female-specific 3D genome dynamics that has both functional and priming effects on the expression of the neuronal genome and a potential to contribute to reproductive hormone-induced brain plasticity and female-specific risk for brain disorders.

## Acknowledgement

This work was supported by the National Institutes of Health: the National Institute of Mental Health under Award Number R01MH123523 (to M.K.) and the National Institute of General Medical Sciences under Award Number R00GM127671 (to M.J.R.). S.C.F. holds the Kim B. and Stephen E. Bepler Professorship in Biology. We would like to thank: Yu Zhang for his assistance with nuclei sorting; Jidong Shan and Cristina Montagna for their assistance with FISH; and Samuel Phillips for his assistance with massively parallel sequencing.

## Author Contributions

M.K. conceived and designed the study; M.J.R. and M.K. directed the project and provided funding; D.R. performed all animal work, immunocytochemistry, tissue preparation for FISH, and assisted with nuclei prep and sorting for Hi-C; M.K. isolated neuronal nuclei, performed the Hi-C assay, and constructed Hi-C libraries; M.S. and M.J.R. processed the Hi-C data, including mapping, quality filtering, and comparison of replicates, as well as the identification and analysis of compartments, CTCF loops, and enhancer-promoter interactions; M.S. and M.J.R. performed reprocessing of the ChIP-seq, ATAC-seq, and RNA-seq data and comparison to Hi-C; A.K. performed gene enrichment pathway analyses; D.R. performed the analysis of FISH data; S.C.F. performed confocal microscopy; D.R., M.S., S.C.F., M.J.R. and M.K. interpreted the data and constructed the figures; M.K. wrote the article with contributions from all co-authors.

## Data and materials availability

Hi-C data are available from the NCBI Gene Expression Omnibus database under accession number GSE172228. ATAC-seq and nuclear RNA-seq data on sorted vHIP neurons from diestrus, proestrus and male groups were previously generated in triplicates (*14*) and are available from the NCBI Gene Expression Omnibus database under accession number GSE114036.

All other data are included in the publication or available from the authors upon request. Correspondence and requests for materials should be addressed to M.K. and M.J.R.

## Supplementary Materials

## Materials and Methods

### Animals

Male and female C57BL/6J mice from Jackson Laboratory arrived at seven weeks of age and were housed in same-sex cages (n = 4-5 per cage). Mice were habituated for two weeks and were kept on a 12:12h light:dark cycle (lights on at 8 a.m.) with *ad libitum* access to food and water. Following habituation, the estrous cycle of female animals was tracked daily in the morning (between 9 a.m and 11 a.m) for two weeks in order to establish a predictive cycling pattern for each female animal and to ensure that only females with regular cycles were included in the study. At 11 weeks of age, male and female animals were sacrificed via cervical dislocation; brains were extracted, and bilateral ventral hippocampi were dissected on ice then flash frozen in liquid nitrogen. For histology experiments, the whole brain was dissected and preserved for cryosectioning (see *Tissue preservation for histology*). Frozen tissue was stored at −80°C before further processing. All animal procedures were approved by the Institutional Animal Care and Use Committee at Fordham University.

### Estrous stage determination

The mouse estrous cycle is typically 4-5 days in duration and contains the following four phases: proestrus, estrus, metestrus, and diestrus. Estrous cycle stage of female animals was determined using vaginal smear cytology, as previously described (*14*). Briefly, smears were collected by filling a disposable transfer pipette with 100μl of distilled water, gently placing the tip of the pipette at the vaginal opening and collecting cells via lavage. The cell-containing water was then applied to a microscope slide and allowed to dry at room temperature for two hours. Once dried, slides were stained with 0.1% crystal violet in distilled water, washed, and then allowed to dry prior to examination with light microscopy. Estrous cycle stage can be determined by the relative quantities of nucleated epithelial cells, cornified epithelial cells, and leukocytes. Establishing the cycling pattern of each female animal allowed us to select days for tissue collection that would maximize the number of females in the proestrus and diestrus phases. Proestus and early diestrus were selected as groups for molecular and histological analysis due to their hormonal profiles mimicking the follicular and luteal phases of the human menstrual cycle, respectively. We have previously confirmed that proestrus represents the high estadiol-low, progesterone phase and early diestrus represents the low estradiol-high progesterone phase (*14*). Phase predictions were confirmed after sacrificing by collecting and analysing post-mortem vaginal smears.

### Nuclei isolation and fluorescence-activated nuclei sorting (FANS) for Hi-C assay

Purification of neuronal nuclei was performed as described previously (*27*) with some modifications to include formaldehyde cross-linking. Briefly, 6 animals per each group - proestrus, diestrus, and males (total n = 18) - were used for Hi-C analysis. From each animal, we used bilaterally dissected ventral hippocampi and pooled brain tissue from two animals for each biological replicate (n = 3 replicates/group). Nuclei preparation and sorting were performed in three batches (n = 3) with each batch having equal group distribution (n = 1 for proestrus, diestrus and males). Brain tissue was dissociated in lysis buffer using a douncer, then incubated in 1% formaldehyde for 10 minutes, followed by quenching with 200 mM glycine for 5 minutes. After washings and filtration through 70 um strainer (Sigma), nuclei preparation continued as described previously (*27*). Total nuclei were extracted using sucrose gradient centrifugation. The nuclei pellet was resuspended in DBPS and incubated, for 45 minutes, with the mouse monoclonal antibody against neuronal nuclear marker NeuN conjugated to AlexaFluor 488 (MAB377X; Millipore, MA). Before sorting, we added DAPI to the incubation mixture and filtered all samples through a 35-μm cell strainer. FANS was performed on a FACSAria instrument (BD Sciences), at the Albert Einstein College of Medicine Flow Cytometry Core Facility. In addition to a sample containing NeuN-AlexaFluor 488 and DAPI stain, three controls were used to set up the gates for sorting: DAPI only; IgG1 isotype control-AlexaFluor 488 and DAPI; and NeuN-AlexaFluor 488 only. We set up the protocol to remove debris, ensure single nuclear sorting (using DAPI), and select the NeuN+ (neuronal) and NeuN-(non-neuronal) nuclei populations (**Fig. S8A**). For each biological replicate, we collected 200,000 NeuN+ (neuronal) nuclei in BSA-precoated tubes filled with 200 μL of DPBS. The purity of sorted single nuclei was confirmed using fluorescence microscopy (**Fig. S8B**).

### Hi-C assay

After FANS, 200,000 neuronal nuclei were pelleted in a centrifuge at 2,850 × *g* for 10 minutes at 4°C, supernatant was removed, and the pellet was frozen in liquid nitrogen and stored at −80°C. Cross-linked, sorted nuclei were then used for the Hi-C assay which was performed using the Arima Hi-C kit (Arima Genomics), according to the manufacturer’s instructions. Briefly, the cross-linked chromatin was digested using a restriction enzyme cocktail; digested ends were filled in and labelled with biotin, followed by the ligation of spatially proximal digested ends. The proximally-ligated DNA was then purified using SPRIselect DNA purification beads (Beckman Coulter) and the first quality control checkpoint was performed to ensure that: a) the sufficient fraction (>15%) of proximally-ligated DNA was labelled with biotin; and b) the output of the Hi-C assay was of the expected size of 2.5-8 kb. The proximally-ligated DNA was then fragmented using a Covaris S2 instrument, targeting the DNA fragment size of 400 bp, which was confirmed using the Agilent Bioanalyzer instrument. This was followed by DNA size selection (200-600 bp) using SPRIselect beads. The size-selected biotinylated fragments were enriched with the Enrichment Beads (provided in the Arima Hi-C kit) and used for Hi-C library preparation. As recommended for low-input Hi-C protocols, the Hi-C library was prepared using the Swift Biosciences Accel 2S Plus DNA Library Kit and Indexing kit reagents, using a modified library preparation protocol in which DNA remains bound to the Enrichment Beads. Following end repair and adapter ligation and before PCR amplification, the second quality control was performed using the KAPA Library Quantification kit (Roche), to determine DNA recovery and number of PCR cycles needed for each library. The library amplification step included 6-7 PCR cycles and was performed using the KAPA Library Amplification kit (Roche). The final Hi-C library was purified using SPRIselect beads and the quality and size of the library (approximately 500 bp) were confirmed using the Agilent Bioanalyzer. The libraries were quantified by Qubit HS DNA kit (Thermofisher Scientific) and the KAPA Library Quantification kit prior to sequencing. 150 bp, paired-end sequencing was performed on a Nova-Seq 6000 instrument at the New York Genome Center. A library pool containing equal amounts of all 9 libraries (n = 3 for proestrus, diestrus, and males) was prepared and loaded on two lanes, yielding close to 200 million reads per library (**Suppl. Table 1)**.

### Hi-C data analysis

Sequenced reads were mapped using BWA (*58*) to the mouse genome build mm10, with a quality filter ≥ 10, and removal of PCR duplicates. Individual replicates were assessed by comparing the distance normalized signal as derived by the formula (observed + 1) / (expected + 1), after which the similarity metric and clustering was derived using DESeq2’s (*59*) sample-to-sample distance function. Replicates were combined with random sampling of reads to have equal levels between samples, and then visualized with Juicebox. Compartments were identified by calculating the eigenvector on the Pearson correlation matrix in 25 kb bins. Differential compartments were identified if the eigenvector differences were higher than 0.42 which corresponded to the differences between quartiles; in essence differential compartments were those that shifted from one quartile to another. CTCF loops were identified using SIP (*29*) at 10 kb resolution using the following parameters: -norm KR –g 1 –mat 2000 -fdr.01. Differential CTCF loops were called by first merging loops from each sample into a master loop list keeping only one entry if loops were called within 25 kb of each other. Then to call strong differential loops, we required a distance-normalized signal >2 and a fold change between samples ≥2.5. Enhancer-Promoter (E-P) interactions were identified by FitHiC2 (*30*) at 10 kb resolution keeping interactions with a q-vaule <.05 and removing any that could be considered (within 20 kb) a CTCF loop instead. We then categorized interactions based on overlap with promoters and non-promoter ATAC-seq peaks as a proxy for enhancers. Differential enhancer-promoter interactions were identified for FitHiC2 interactions with a distance-normalized signal >2 and a fold change between samples ≥2.5. Comparison to Hi-C and H3K27me3 in ES, NPC, and CN was done by reprocessing data from GEO accession GSE96107 (*60*) and from ENCODE accession ENCSR059MBO. Metaplots of these and other interactions were created using SIPMeta (*29*).

### ATAC-seq and RNA-seq data

ATAC-seq and nuclear RNA-seq data on sorted vHIP neurons from diestrus, proestrus and male groups were previously generated in triplicates (*14*) and are available from the NCBI Gene Expression Omnibus database under accession number GSE114036.

### Integration of Hi-C, ATAC-seq, and RNA-seq data

ATAC-seq and nuclear RNA-seq previously processed and mapped to mm10 (*14*) was used in conjunction with ngs.plot (*61*) to create average expression profiles as TSSs. Motifs were obtained by meme-chip (*60*). Genome browser tracks display bins per million mapped reads (BPM) normalized signal and were visualized alongside arc-plots of interactions using the WashU Epigenome Browser (*62*). Gene expression was evaluated by Stringtie (*63*) to provide transcripts per million (TPM) normalized values. X escapee genes were identified as those with TPM ≥ 1 in females and ≥1.5 fold difference to males. Gene ontology and pathway analyses were performed by EnrichR (*64*) and the Ingenuity Pathway Analysis (IPA) software (https://www.qiagenbioinformatics.com/products/ingenuitypathway-analysis). Comparison of expression in other tissues was done using the Human Protein Atlas database (*31*) and for disease models using the Mouse neurological disorders RNA-seq portal (*39*).

### Tissue preservation for histology

Following whole brain isolation, brains were washed with ice cold 0.1 M PBS and fixed in 4% PFA in 0.1M PBS at 4°C for 24h. After fixation, brains were rinsed in cold 0.1 M PBS and underwent sucrose preservation, which involved placing the brains in solutions containing 15% then 30% sucrose dissolved in 0.1 M PBS at 4°C for 24h and 48h, respectively. Brains were then frozen in dry ice-cooled hexane and stored at −80°C until sectioning. Cryosectioning was performed by embedding brains in optimal cutting temperature compound (OCT) and cutting serial sections on a rotary cryostat (Leica CM1850, Leica Biosystems GmBH). 5-10μm coronal sections containing the ventral hippocampus were collected on Super Frost Ultra Plus slides (Fisher Scientific) and were processed for either fluorescent in-situ hybridization (FISH) or immunofluorescence microscopy.

### Fluorescence in-situ hybridization (FISH)

Cryopreserved brain sections from n = 3 animals/group were sent to Albert Einstein College of Medicine Cytogenetics Core Facility for FISH. Briefly, four bacterial artificial chromosome (BAC) clones corresponding to the regions of interest were obtained from the BACPAC Resources Center (Children’s Hospital Oakland Research Institute). DNA was isolated from bacterial clones, labeled with fluorophores, and hybridized to tissue sections, as previously described (*65*). Images were acquired using a Zeiss Axiovert 200 inverted microscope (Carl Zeiss MicroImaging, Inc.) using fluorophore-specific filters. The targeted regions were as follows (mm10): Probe 1 - chrX: chrX:153798639-153942772 (BAC RP24-305G7, Red label); Probe 2 - chrX: 143115199-143348205 (BAC RP24-88L14, Aqua label); Probe 3 - chrX: 148937622- 149119722 (BAC RP24-295N13, FITC label); Probe 4 - chr1: 133514089-133681697 (BAC RP24-287A12, Yellow label). Data analysis of FISH images was performed using ImageJ (public domain software from the National Institutes of Health; http://imagej.nih.gov/ij/). For each X chromosome the distance between the center of the red and blue signals (RB) and the red and green signals (RG) was measured. The difference between these measurements (RB-RG) was calculated for each nucleus in each group and data analysis was performed using one-way ANOVA with Tukey as a posthoc test. The analysis was restricted to X chromosomes positive for all three probe signals (n=116-163/group).

### Immunofluorescence microscopy

Slides with cryopreserved brain sections were rehydrated in PBS and underwent blocking in 0.25% Triton X-100 and 5% BSA in PBS for 1 hour at room temperature. Following blocking, the slides were incubated at 4°C for 24 hours with the following primary antibodies: (I) rabbit polyclonal anti-ERα antibody (1:2,000; Sigma-Aldrich, 06-935) (II) mouse monoclonal anti-NeuN conjugated to AlexaFluor-488 (1:500; Millipore, MAB377X). Slides were subsequently incubated in the dark for 2 hours at room temperature with secondary antibody donkey anti-rabbit IgG conjugated to AlexaFluor-594 (1:250; Invitrogen, A-21207). After washing with PBS, the slides were counterstained with DAPI (1:1,000) and mounted with Mowiol 4-88 mounting medium (Sigma-Aldrich). Imaging was performed using a Leica TCS SP8 confocal microscopy system (Leica Microsystems CMS GmbH).

---

**Figure S1.**
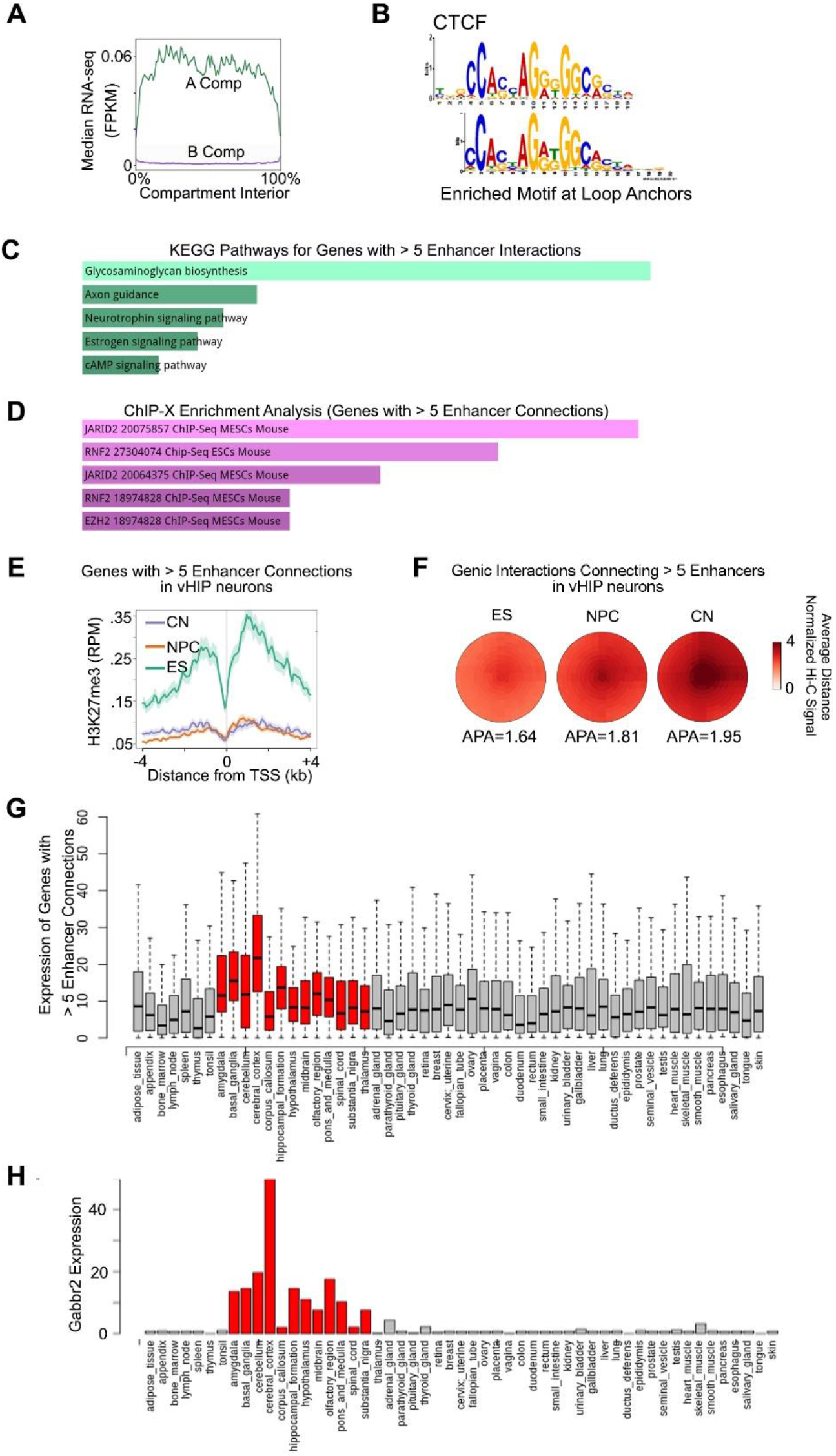
Multi-level 3D genome organization in vHIP neurons. The A chromosomal compartment is associated with active transcription (strong RNA-seq signal) while the B compartment is associated with transcriptionally inactive genes (**A**). Anchor regions of CTCF loops called by SIP are enriched for the CTCF motif (**B**). Genes with more than 5 enhancer-promoter (E-P) interactions in vHIP neurons are involved in neuronal function and hormone signaling as shown by the KEGG pathway analysis (**C**) and are enriched for binding of repressive polycomb proteins in mouse embryonic stem cells (ESCs) as shown by ChIP-X enrichment analysis (**D**). The same genes are enriched for the repressive histone modification H3K27me3 in embryonic stem (ES) cells, but not in neural progenitor cells (NPC) or cortical neurons (CN, **E**) and the observed E-P interactions become stronger over the course of neuronal differentiation (**F**). The genes with over 5 E-P interactions in vHIP neurons are primarily expressed in the central nervous system (**G**) including *Gabbr2*, encoding a subunit of the GABA-B receptor (**H**). Data by Bonev et al. (*18*) were used to examine H3K27me3 levels and E-P interactions in our multi-enhancer genes during neuronal differentiation. Data by Uhlen et al. (*31*) were used to test the expression of these genes across tissues.

**Figure S2.**
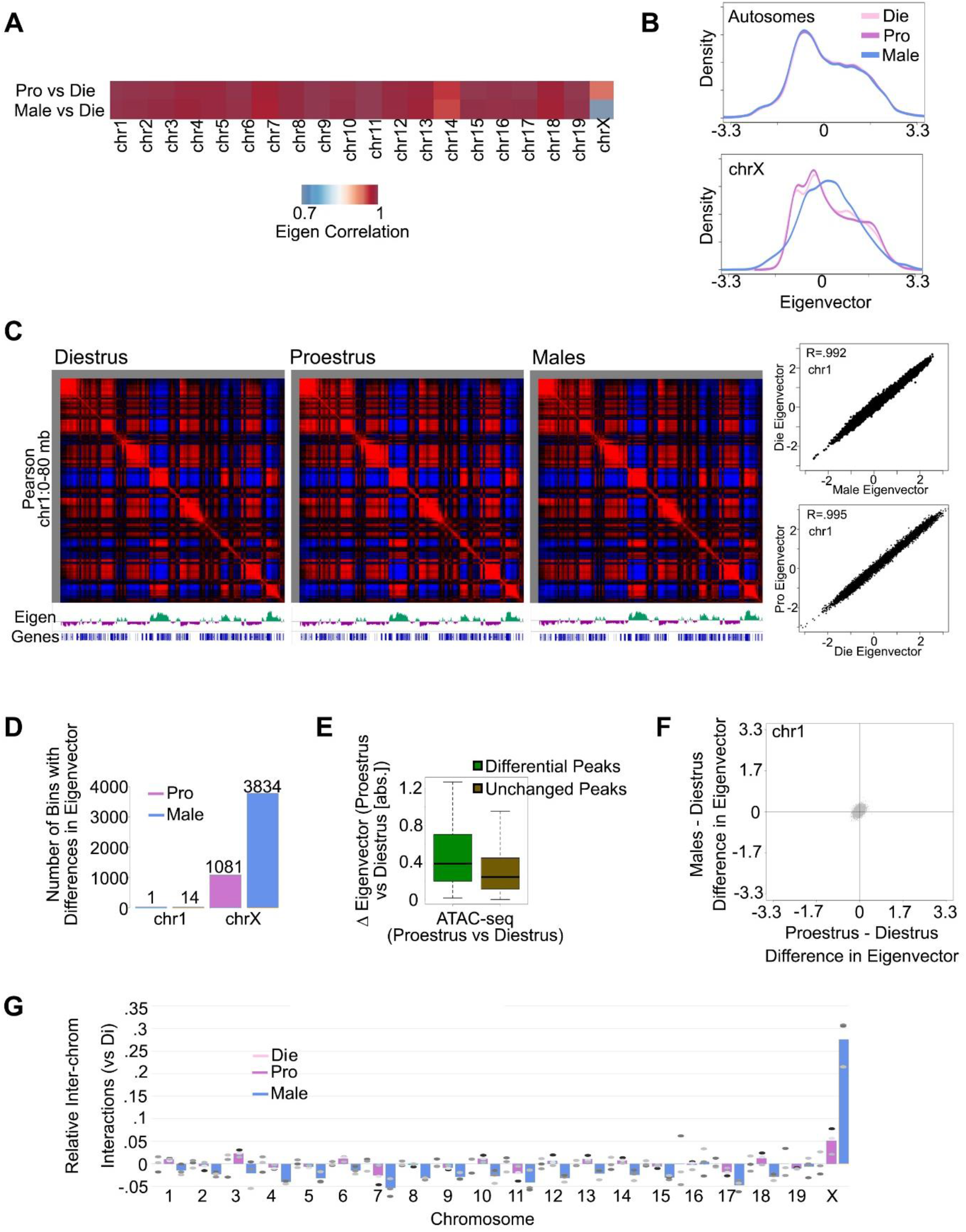
Compartmental organization in vHIP neurons across sex and the estrous cycle. Sex- and estrous cycle-dependent compartmental changes, displayed by eigenvector heatmaps (**A**) and density plots (**B**) are primarily observed on the X chromosome. A correlational Hi-C matrix with eigenvector signal (Eigen) for an 80 Mb-region of the chromosome 1 (left) shows a high correlation (right) of compartmental signal in Die-Male and Die-Pro comparisons (**C**). The number of 25-kb bins with differences in eigenvector across-sex and within-females for the X chromosome and chromosome 1 (**D**). X chromosome bins with differences in eigenvector within-females also display differential chromatin accessibility (**E**). No association was observed between Pro-Die and Die-Male compartmental changes on the chromosome 1 (**F**). When each separate biological replicate was examined (see differentially colored ellipses), proestrus and males display a greater number of inter-chromosomal interactions on the X-chromosome, but not on autosomes, compared to diestrus (**G**).

**Figure S3.**
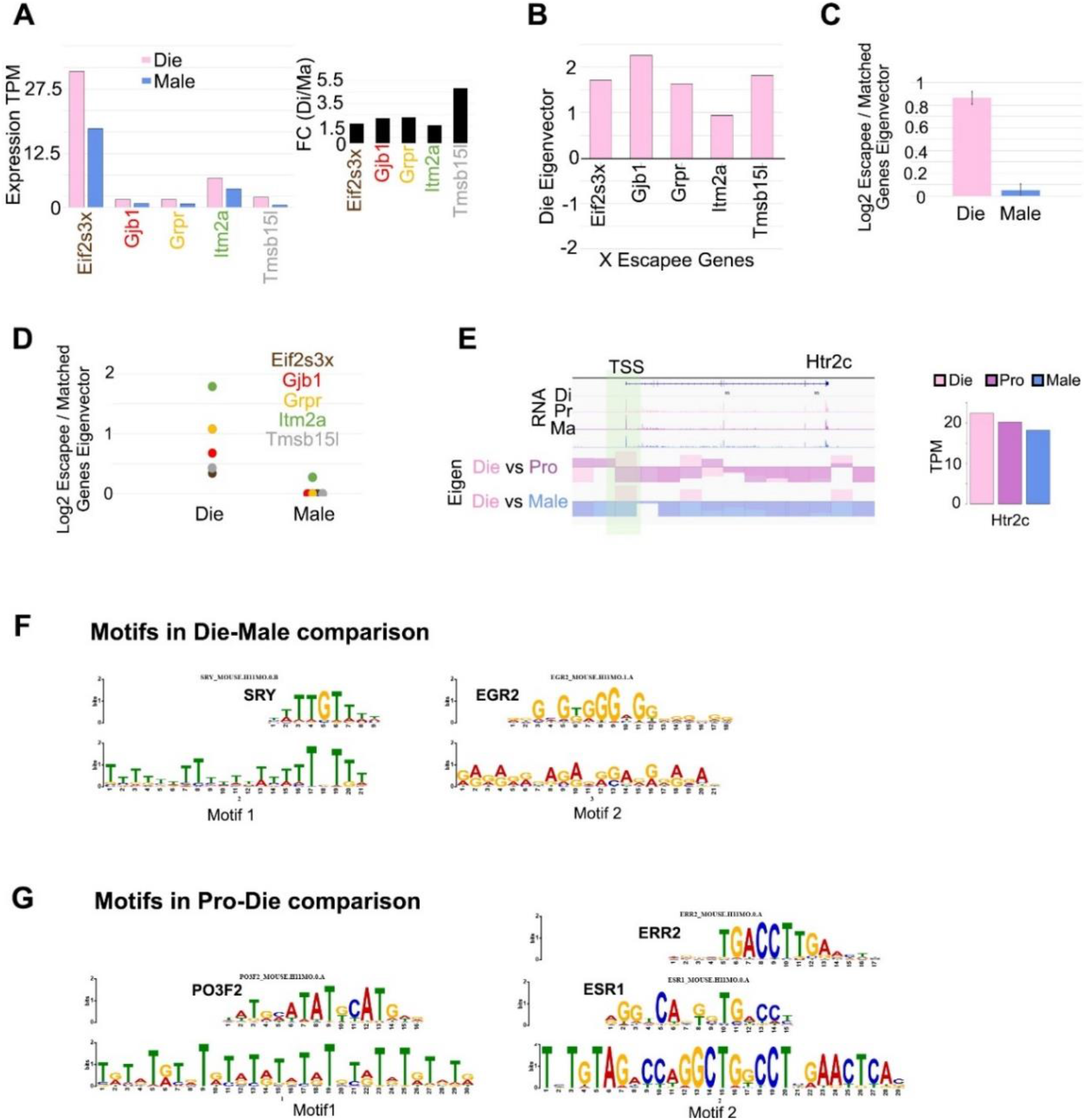
X chromosome compartmental differences are associated with sex-specific gene expression and motif enrichment. Expression levels (left) and expression fold-change (right) of five X-linked genes in the Die-Male comparison (**A**), along with their A compartmental profiles in diestrus (**B**) indicate these genes undergo X-escape. Relative to genes with similar expression levels, expression of these escapees is associated with a higher eigenvector signal in diestrus compared to males on average (**C**) and by individual escapee (**D**). Eigenvector tracks for *Htr2c* (left) indicate differential compartmental profiles across the estrous cycle and sex that overlap the transcription start site (TSS) and correlate with altered expression levels (right, **E**). Note that we previously identified *Htr2c* as a variable X escapee in adult vHIP neurons, which is differentially expressed in the Die-Male but not in the Pro-Male comparison (*14*). Motif analyses of genomic regions with differential compartment signals between groups show enrichment for Sry and Egr2 binding sites in the Die-Male comparison (**F**), and enrichment for Pou3f2, ERα, and ERR2 binding sites in the Pro-Die comparison (**G**).

**Figure S4.**
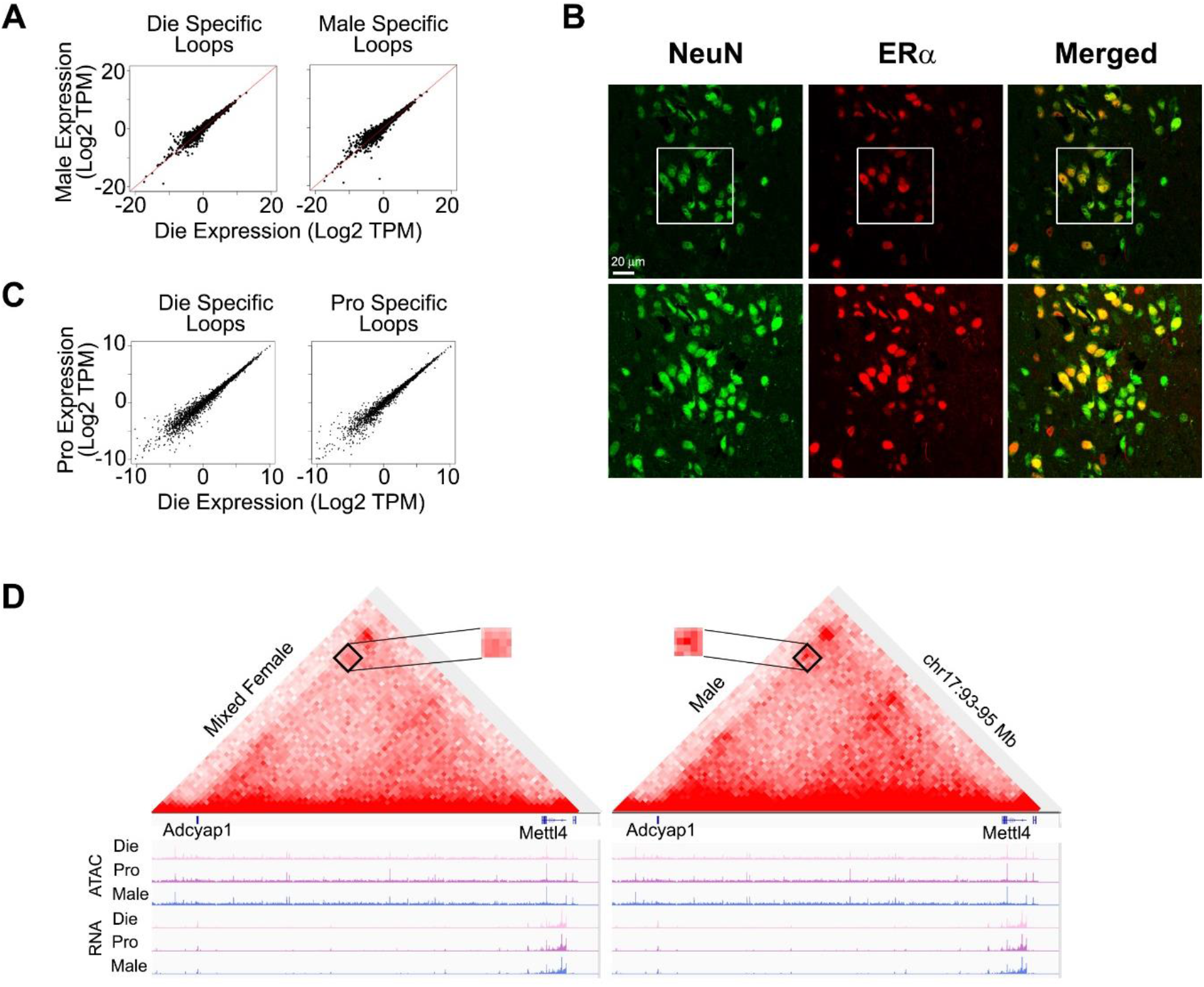
CTCF loops in vHIP neurons across sex and the estrous cycle. On a global level, gene expression in diestrus and males is similar whether the gene is within a diestrus- or male-specific loop (**A**). Confocal microscopy images of vHIP tissue sections stained for NeuN, a marker for neuronal nuclei, and ERα, with an overlap indicating ERα is present in vHIP neuronal nuclei, consistent with its possible role as a loop organizer in these cells (**B**). The same field is shown as x-y section (upper row) and the maximum intensity projection (lower row); the smaller, boxed region was shown in the main Figure 4E. While the image shown depicts a proestrus sample, nuclear ERα was observed in NeuN+ cells of all three groups (**B**). On a global level, gene expression in proestrus and diestrus is similar whether the gene is within a proestrus- or diestrus-specific loop (**C**). A specific, 2Mb-loop connecting *Adcyap1* and an upstream region of *Mettl4* found to be stronger in proestrus than in diestrus (**Fig. 4F**), is identified as a differential loop in merged-females compared to males, with a weaker signal in females, emphasizing a loss of specificity when the female groups are merged (**D**).

**Figure S5.**
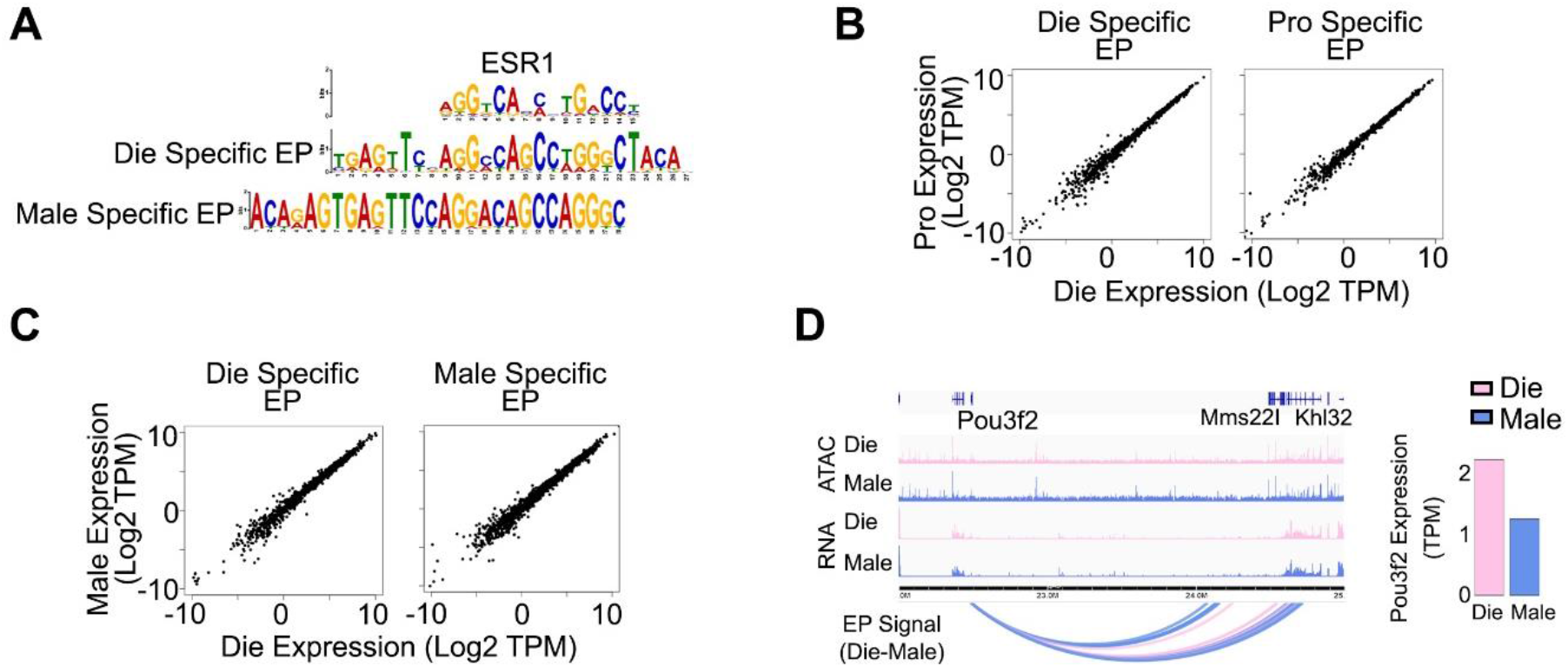
E-P interactions in vHIP neurons across sex and the estrous cycle. Genomic regions with differential enhancer-promoter (E-P) interactions in the Die-Male comparison are enriched with ERα binding sites (**A**). Gene expression within-females is similar whether the gene has Die- or Pro-specific E-P interactions (**B**), and gene expression across sex is similar whether the gene has Die- or Male-specific E-P interactions (**C**). Pou3f2 exhibits differential E-P interactions in the Die-Male comparison that are associated with differential gene expression (**D**). IGV tracks show merged ATAC-seq and RNA-seq data for diestrus (Die), and males (Male), and the bar graph (on the right) shows Pou3f2 gene expression. All data were derived from 3 biological replicates for each group.

**Figure S6.**
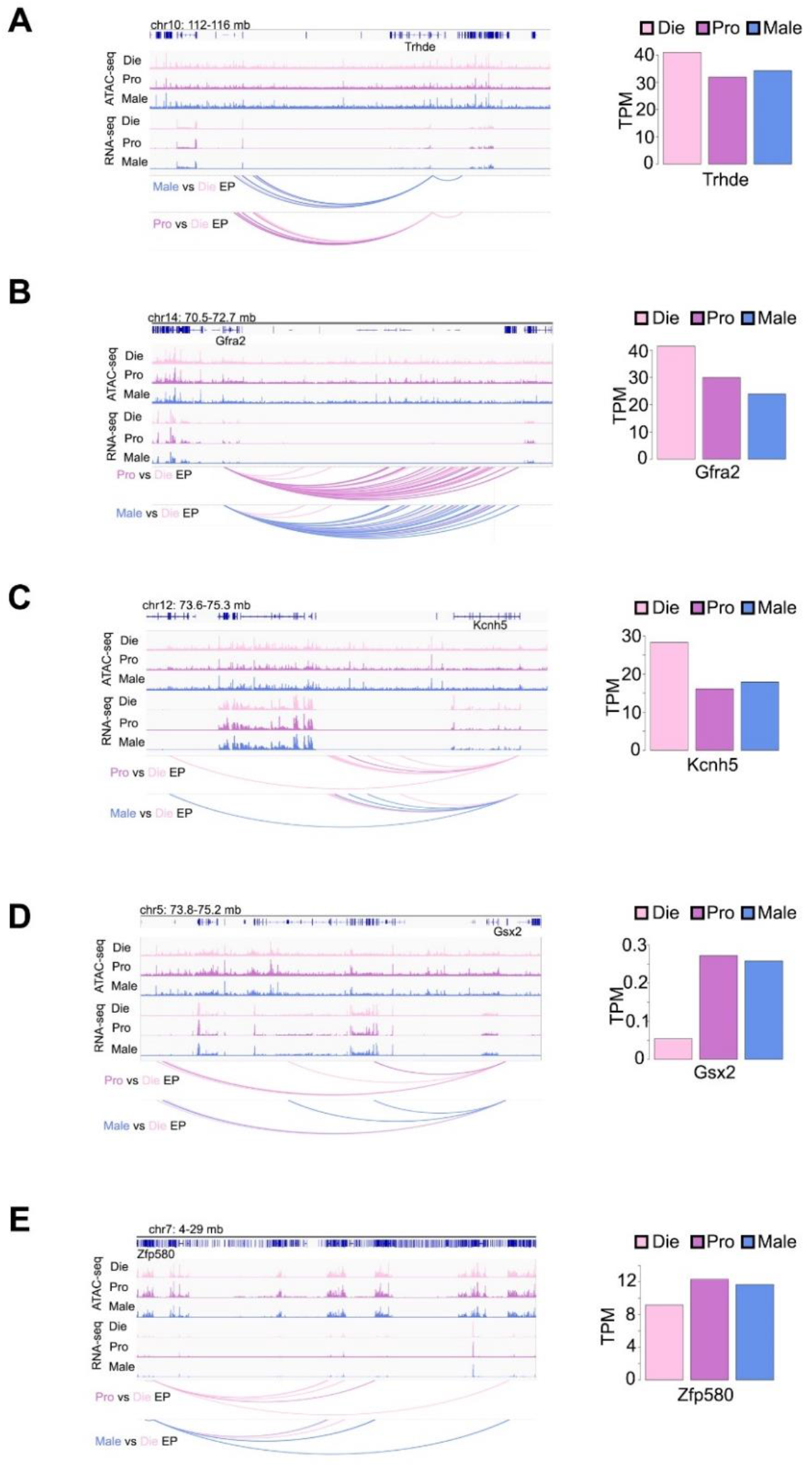
Sex-specific E-P interactions are associated with gene expression in vHIP neurons. Example genes showing differential E-P interaction profiles and differential gene expression dependent on sex and estrous cycle stage including *Trhde* (**A**), *Gfra2* (**B**), *Kcnh5* (**C**), *Gsx2* (**D**), and *Zfp580* (**E**). IGV tracks show merged ATAC-seq and RNA-seq data for diestrus (Die), proestrus (Pro), and males (Male) derived from 3 biological replicates for each group. Bar graphs (on the right) show gene expression for each gene. Die, pink; Pro, purple; Male, blue.

**Figure S7.**
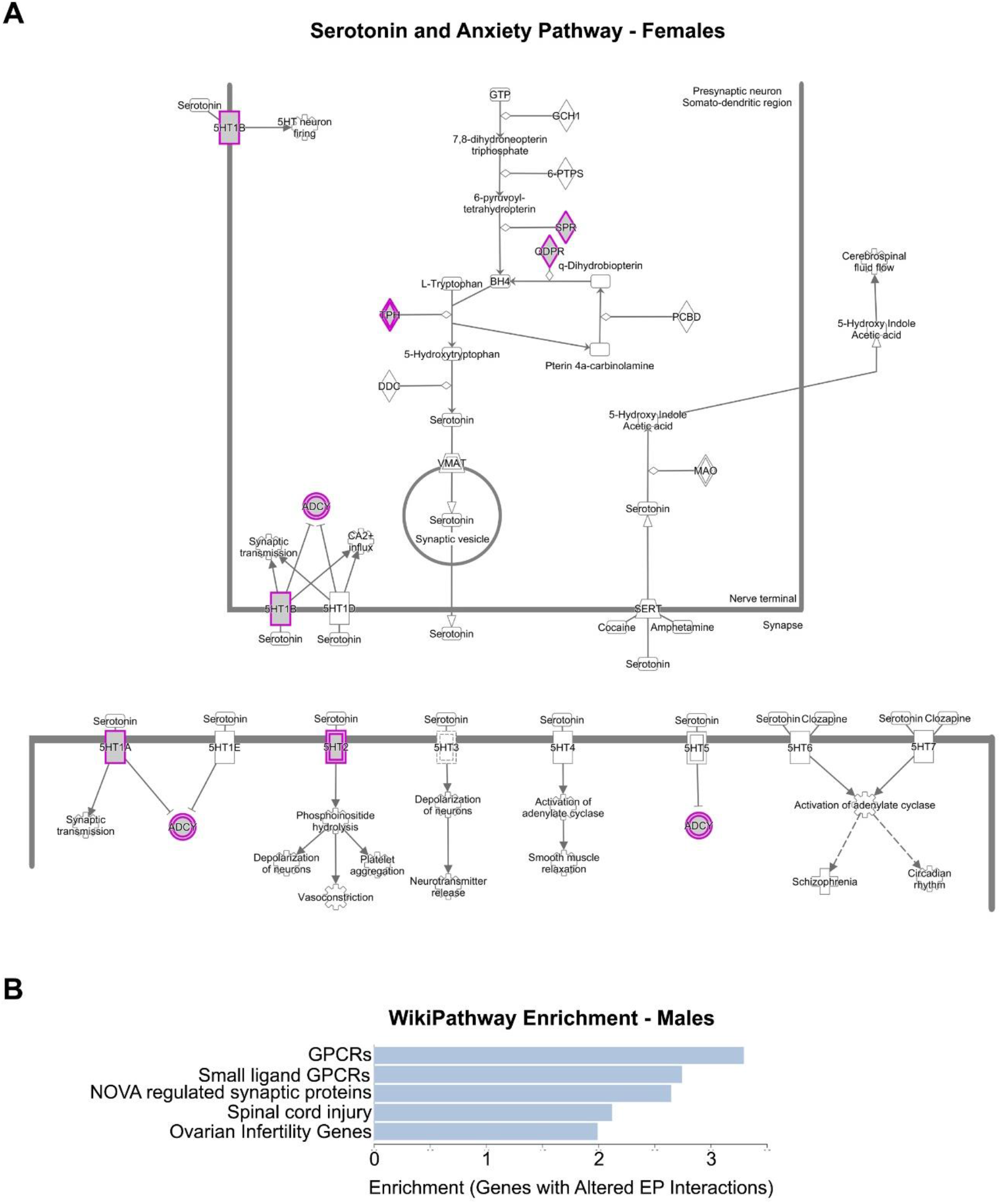
Sex-specific pathway enrichment of differential E-P interactions. (**A**) WikiPathway schematic of the Serotonin and Anxiety pathway. Proteins with purple borders are encoded by genes with differential E-P interactions within-females across the estrous cycle. (**B**) In the Die-Male comparison, top enriched pathways of differential E-P interactions are involved in G protein-coupled receptor (GPCR) signaling.

**Figure S8.**
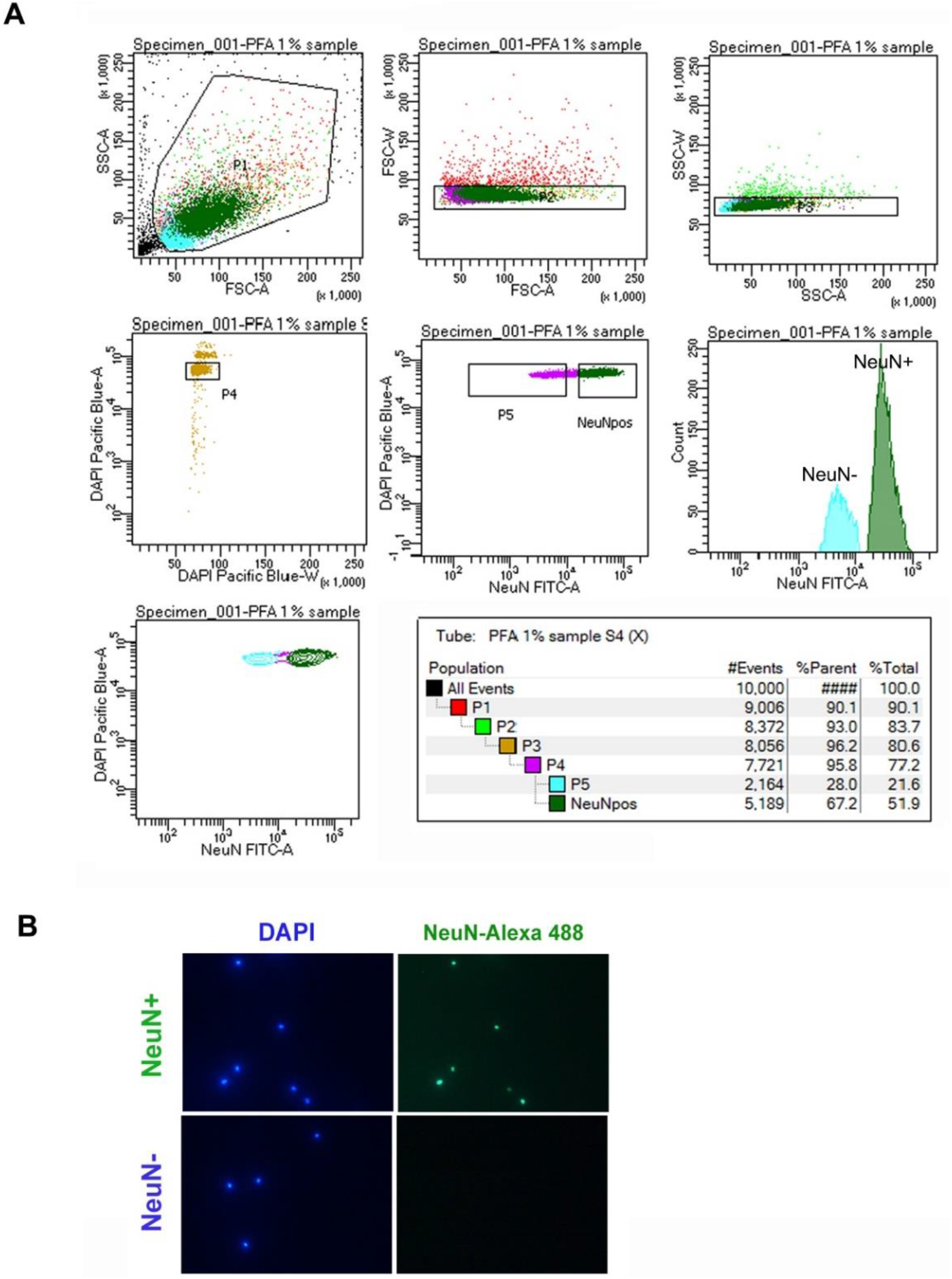
Separation of neuronal nuclei using fluorescence-activated nuclei sorting (FANS). A representative FANS report shows the gating strategy that was used to: 1) separate nuclei from debris (P1-P3); 2) ensure the sorting of single nuclei using the DAPI signal (P4); and 3) select the NeuN+ neuronal nuclei (P6) from the NeuN- non-neuronal nuclei (P5) (**A**). Representative immunofluorescence microscopy images attained after sorting demonstrate that sorting separates NeuN+ from NeuN- nuclei and results in a single-nuclei suspension (**B**).

**Supplementary Tables (separate files)**

**Table S1:** Hi-C data basic information (<u>uploaded as a separate excel file</u>)

**Table S2:** List of genes with multiple E-P interactions (<u>uploaded as a separate excel file</u>)

Suppl. Table S2a: Genes with over 5 enhancers;

Suppl. Table S2b: Genes with 3-5 enhancers.

**Table S3:** List of genes associated with differential loops and E-P interactions across the estrous cycle (<u>uploaded as a separate excel file</u>)

Suppl. Table S3a: Genes Closest to Die-Specific Loop Anchors;

Suppl. Table S3b: Genes Closest to Pro-Specific Loop Anchors;

Suppl. Table S3c: Genes at Die-Specific EP Anchors;

Suppl. Table S3d: Genes at Pro-Specific EP Anchors.

